# Structural and Dynamical Analyses of Apo and Cap-binding eIF4E: An *in silico* Study

**DOI:** 10.1101/2024.05.18.594816

**Authors:** Karin Hasegawa, Justin Burzachiello, Liam Connolly, Georgios Kementzidis, Ziyuan Niu, Evangelos Papadopoulos, Bertal Huseyin Aktas, Yuefan Deng

## Abstract

Molecular dynamics (MD) simulations were utilized to investigate the dynamic behavior of eukaryotic translation initiation factor 4E (eIF4E) in its apo (ligand-free) and Cap-bound (complexed with mRNA Cap) forms. Specifically, we focused on the rearrangement of critical eIF4E residues, including W102, E103, W56, and key charged residues within the alpha helix, such as lysine and arginine. Three independent 1 *µs* MD simulations were performed for four eIF4E structures based on an available Cap-bound eIF4E crystal structure (PDB: 4TPW) to sample protein conformations and explore metastable states. Metastable states were successfully sampled within the 1 *µs* trajectories for all four structures; however, 2D-root-mean-square deviation (RMSD) and free-energy landscape (FEL) analyses indicate that additional trajectory data may be needed to fully explore the conformational space. Notably, the clearest state transitions were observed in the Cap-A simulations. Our data suggests that previously reported allosteric site eIF4E inhibitor 4EGI-1 stabilized the ^7^mGTP-eIF4E interaction by binding to a site dorsal from the eIF4G binding site, facilitating stable protein states even after ligand removal and promoting cleaner state transitions in simulations. Root-mean-square fluctuation (RMSF) and distance analyses aligned consistently with experimental results, affirming the validity of our MD simulation system. Observations on apo and Cap eIF4E structures revealed structural changes in one of the alpha helices (*i.e.,* H3), particularly the transition from alpha helix to loop upon Cap binding. This transition was also observed during the cap to apo transition, where residues E105-R109 underwent structural rearrangements from loop to alpha helix. These findings support the strategy of designing agents targeting critical Cap-binding residues (W56, W102, E103) and H3 lysine residues (K106, K108) to disrupt the ^7^mGTP-eIF4E interaction, potentially informing future drug design efforts targeting this oncogenic protein. Future work will focus on studying the apo to Cap transition using available eIF4E apo structures, such as PDB entry 2GPQ, to deepen our understanding of eIF4E’s metastable states during Cap binding.

## Introduction

Mammalian translation initiation plays a critical role in regulating gene specific expression, *i.e.,* deciding which proteins encoded by which genes will be produced at what frequency. The initiation stage of protein synthesis involves assembly of a complex that contains mRNA, eukaryotic translation initiation factors (eIFs), and small ribosome subunit (40S); the machine that translate mRNA into proteins. This complex is called the 48S preinitiation complex, which will identify the first amino acid codon (usually methionine) and recruit 60S ribosome subunit to assemble translation competent ribosome. eIF4E is the critical protein that bring the mRNA to the 48S complex. This is because eIF4E binds the mRNA on one hand and eIF4G on the other hand. eIF4G is the large scaffolding protein that binds to not only eIF4E and myriad other translation initiation factors but also 40S ribosome subunit thus delivering mRNA to the translation machinery. eIF4E does not simply bind all mRNAs with similar affinity but has the unique ability to select which mRNAs it will bind based on mRNAs’ primary sequence and/or secondary structures. Importantly, the repertoire of mRNAs recruited to translation machinery is highly dependent on the cellular level and activity eIF4E. Indeed, changing the level of its eIF4E or the activity of eIF4E causes a major shift in the composition of mRNAs recruited for translation. When translationally active eIF4E is abundant, all mRNAs are translated, albeit with varying efficiency. However, when eIF4E expression or activity is restricted, it favors translation of mRNAs required for maintaining cell and tissue homeostasis.^1^ Therefore, abundance of eIF4E is essential for translation of mRNAs coding for proteins that play critical roles in regulating cellular functions such as proliferation, malignant transformation, ^2–7^ mitochondrial dynamics,^8,9^ learning and memory,^10–14^ and aging. ^15,16^ Consistently, studies that increased or reduced eIF4E expression and/or its cellular inhibitors and activators demonstrated that non-physiologic activation of eIF4E significantly contributes to pathophysiology of many human disorders including cancer,^17,18^ diabetes,^19,20^ autism related learning deficits,^21–23^ Parkinson’s,^24^ Huntington’s^25^ and Alzheimer’s diseases^26^ and play a causative role in fragile-X mental retardation syndrome.^27,28^ These studies indicate that reducing eIF4E expression/activity may be beneficial for the management of an array of human disorders. eIF4E/mRNA interactions are mediated by ^7^mGTPppp mRNA Cap binding to the so-called Cap binding pocket. Extensive studies in the last three decades to find inhibitors of mRNA Cap binding to eIF4E have not succeeded in developing any druggable molecule. Specifically, despite the advantage of having X-ray crystal and nuclear magnetic resonance (NMR) structure of eIF4E, numerous classical structure-based drug design (SBDD) approaches have not succeeded in developing a clinical candidate^29–33^ because displacing a bound mRNA from eIF4E is nearly impossible task for a drug that must cross the cell-membrane. Cap binding pocket does not pre-exist but is induced by mRNA Cap through extensive re-arrangement of critical eIF4E residues, namely W102, E103 and W56. Not only the rearrangement of these residues into a pocket is induced by the mRNA-Cap but also formation of this pocket is essential for interactions. Furthermore Cap binding causes dissolution of heli× 105-109 (*i.e.,* H3)^34^ into a loop structure. Once the base of ^7^mGTPppp interacts with W56, W102, and E103, three negatively charged phosphates of the Cap as well as at least two phosphates of the mRNA penultimate nucleotides make extensive ionic contacts with many of lysine and arginine residues of eIF4E, creating an interaction strong enough to withstand nearly any potential drug that can cross the cell membrane. This is because the drugs that cross the cell membrane cannot have negative charge to compete against binding of Cap and RNA phosphates to eIF4E lysine and arginine residues.

These findings indicate that an approach of restraining the movements of W102, E103, W56 and/or dissolution of H3 by developing bi-valent agents that will form a bridge between the aforementioned residues and a stable secondary structure of eIF4E to prevent the formation of mRNAs Cap binding pocket, therefore, most likely blocking the interaction of the Cap with eIF4E.

W102, E103, and W56, the key residues that form the ^7^mGTP mRNA Cap binding pocket, reside on unstructured flexible loops in apo-eIF4E. While apparently disordered on X-ray crystal and NMR structure, these loops most likely form multiple metastable structures; otherwise, they will destabilize the overall structure of eIF4E. We currently have no knowledge of these metastable cryptic pockets/structures. Similarly, interactions of ^7^mGTPppp with W102, E102, and W56 and formation of mRNA Cap binding site must proceed through intermediate (metastable) states, which are currently not known. The detailed analysis of the high-resolution structure of these transition states, both temporally and spatially, will significantly aid in the advancement of innovative methodologies for crafting new chemical entities (NCEs). These NCEs should hinder the formation of the eIF4E Cap binding site, consequently restricting translation initiation. To fill the aforementioned gap, in this work we will use the existing crystal structures of eIF4E-Cap complex (PDB: 4TPW) to study how the apo-eIF4E transitions through intermediate states from the apo form to the Cap-bound form, and vise versa. Molecular dynamics (MD) simulations of the apo and Cap forms will be used to collect the protein and the complex trajectories, and to give deep insights on the re-arrangement of critical eIF4E residues and the secondary structure of *α*-helix containing key residues such as lysine and arginine residues. The observations made through this study may be utilized for developing inhibitors targeting this oncogenic protein in future drug designs.

## Materials and Methods

### Initial Structures

Two crystal structures of eIF4E in complex (PDB: 4TPW),^35^ one with ^7^mGTPppp (7-methylguanosine triphosphate) Cap analogue (labeled as Chain B) and the other one with both Cap and 4EGI-1 (labeled as Chain A), were sourced from the Protein Data Bank. 4TPW Chain A contained a total of 185 amino acid residues starting from HIS33 to VAL217, and 4TPW Chain B contained a total of 183 amino acid residues starting from ILE35 to VAL217. To prepare the eIF4E structures both with and without the Cap for two structures available in 4TPW, Cap and 4EGI-1 were manually removed from the Chain A structure and Cap was removed from the Chain B structure to generate each alternate uncomplexed structure of eIF4E that we term the “apo” structures. Similarly, 4EGI-1 was manually removed from the Chain A structure and no further modification was made for the Chain B structure to prepare the “Cap” structures. In summary, we have prepared four different structures for eIF4E: a 4EGI-1 removed 4TPW Chain A structure that we refer as “Cap-A”, an original 4TPW Chain B structure that we refer as “Cap-B”, an uncomplexed 4TPW Chain A structure that we refer as “Apo-A”, and an uncomplexed 4TPW Chain B structure that we refer as “Apo-B”. These four prepared structures are presented in Figure 1. We generated the initial topology using the CHARMM36 force field^36,37^ within GROMACS 2020.2. ^38–41^ Each eIF4E structure was positioned at the center of a cubic simulation box for both “apo” structures (*i.e.,* “Apo-A” and “Apo-B”) and for “Cap” structures (*i.e.,* “Cap-A” and “Cap-B”), ensuring a minimum 1 nm clearance between the component and each side of the box. These structures were solvated with TIP3P water.^42^ The net charge were +3 across all the systems, and to neutralize the systems three chloride ions were added. Periodic boundary conditions were applied in all three dimensions.

**Figure 1:**
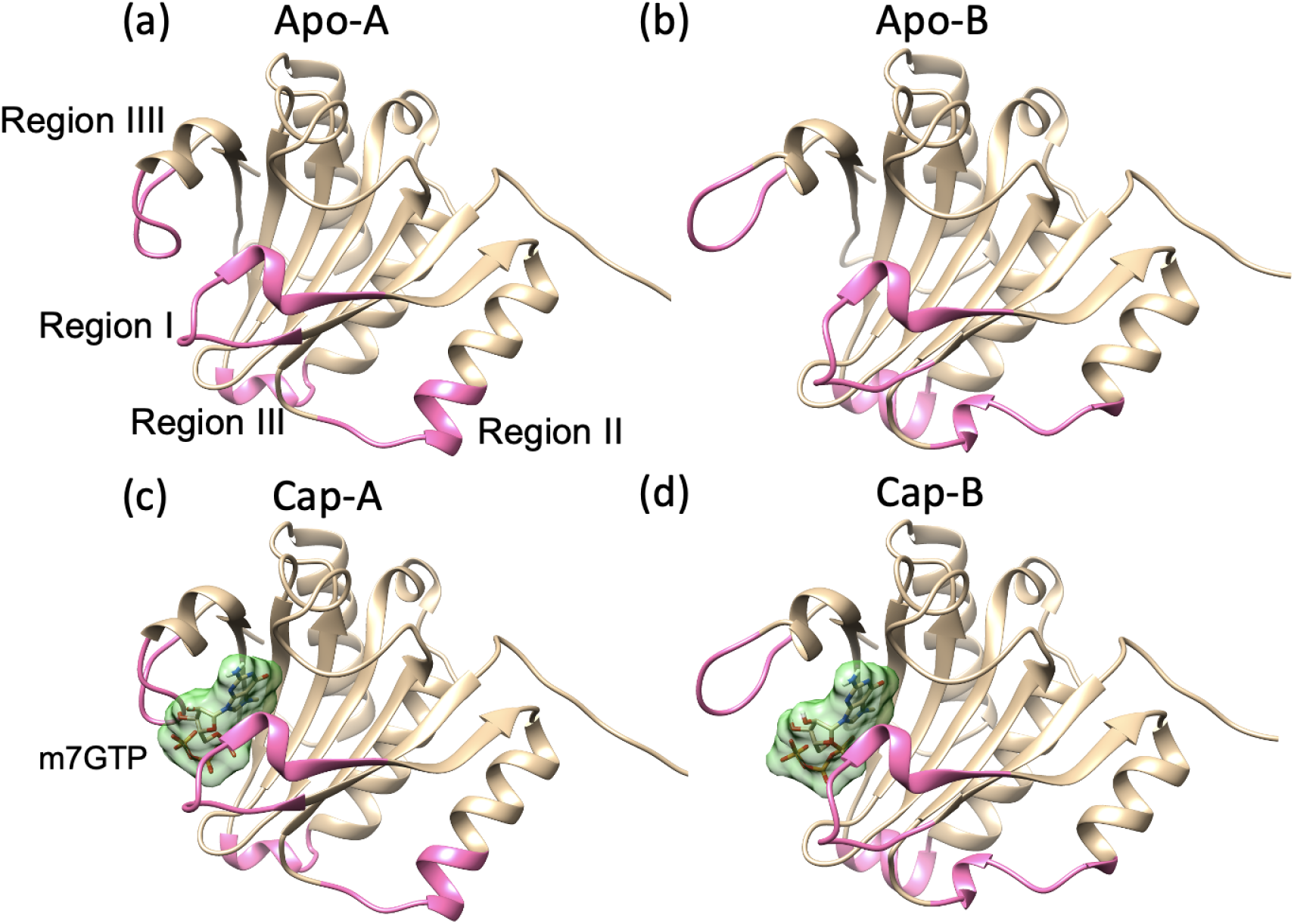
(a) Apo-A, (b) Apo-B, (c) Cap-A, (d) Cap-B. eIF4E is represented as a tan cartoon. Four regions (Region 1: K49-R61, Region 2: H78-L85, Region 3: K119-D125 and Region 4: K206-K212) that showed a large RMSD difference in 4TPW Chain A and 4TPW Chain B structures are colored in hot pink. ^7^mGTP is shown in tan stick with solid lime green surface. All the surfaces are shown in 0.7 transparency for better visibility.

### Molecular Dynamics Simulations

Molecular dynamics (MD) simulations were performed in GROMACS 2020.2^38–41^ with CHARMM36 force field.^36,37^ We initiated energy minimization to the solvated system using the steepest descent algorithm, setting the maximum number of steps to 100,000 with a step size of 0.002 fs and a maximum force threshold of 1000 kJ/mol/nm. For buffered neighbor searching, the Verlet cutoff-scheme was employed. The neighbor list was determined using the grid method and updated at each step. For electrostatic interactions, the particle mesh Ewald method (PME) was used, with a 1 nm cutoff. The same cutoff range was applied for short-range Van der Waals interactions. Following energy minimization, temperature coupling was performed at 310 K using V-rescale algorithms.^43^ Temperature equilibration took place over 100 ps to 150 ps until each system reached the specified temperature. Pressure coupling was performed at 1 atm for 100 ps to 150 ps using the Parrinello-Rahman method.^44^ Throughout both temperature and pressure coupling stages, position restraints of 1000 kJ was imposed to eIF4E for apo-eIF4E systems and to both eIF4E and the Cap for Cap-eIF4E systems. H-bonds were constrained using the LINCS algorithm. Once temperature and pressure converged to the target values, position restraints were fully released for eIF4E in apo-eIF4E systems and for both Cap and eIF4E in Cap-eIF4E systems. Subsequently, 1 *µs* production run was conducted in the NPT ensemble for all four systems. The whole process was repeated three times to prepare the triplicate runs for each structure. Each output row trajectory was then post-processed to only contain 1 frame/ns to reduce the data size for the subsequent analysis steps, resulting in approximately 1000 frames per trajectory after the process. Furthermore, the amino acid residues through LYS36 in eIF4E N-terminus were all ignored from the protein structures in all the analyses except for the RMSF calculations in order to avoid the large flexibility of the domain skewing the data analysis results.

## Results and Discussion

### Global Conformational Change of eIF4E

To identify and collect the ensembles of both apo and Cap structures, 2D RMSD and Free Energy Landscape (FEL) were calculated for all the collected trajectories using MDAnalysis tool^45,46^ and gmx sham command from gromacs, respectively. 2D RMSD, also often referred as all-to-all RMSD, is the plot based on taking the RMSD of each snapshot of an interest in the trajectory with respect to all other snapshots. It is commonly used as a similarity measure of a protein structure at different simulation time points, and a dark square appears in such a plot when the protein structure does not change much during some length of time interval while yellow regions appear when the protein structure has changed significantly. For the 2D RMSD calculation, for each structure we concatenated the trajectory from all three runs into one trajectory which resulted in a 3000 ns long trajectory (*i.e.,* 0-1000 ns, 1000-2000 ns, and 2000-3000 ns correspond to Run 1, Run 2 and Run 3, respectively). Subsequently, each protein trajectory was aligned at its *α* carbon atoms, and then RMSD values were calculated. The resulting 2D RMSD for each structure is presented in Figure 2. For FEL, gmx sham was used to combine the data points of the reaction coordinates of the principle components with the largest motions. FEL is given in kJ/mol and calculated based on the concatenated trajectory (*i.e.,* 2400 ns) of each run’s truncated trajectory (*i.e.,* 200 ns - 1000 ns) where the first 200ns was discarded for the system equilibration. For the calculation, first a covariance matrix was produced based on the trajectory, and the eigenvalues and the corresponding eigenvectors were calculated. The first two largest eigenvalues and the corresponding eigenvectors are then extracted and used to calculate and generate the Gibbs free energy landscape based on the Boltzmann ensemble. The output FEL is presented in Figure 3. The regions in FEL that have exhibited low free energies were highlighted with red rectangular boxes and the corresponding states were also highlighted with the same red boxes in the 2D RMSD. For each eIF4E structure, those highlighted low energy states were named differently so that the states can be distinguished one from another. The assigned state names are displayed right next to each red boxes on both FEL and 2D RMSD plots. The average structures for the low energy states are also presented in Picture A of Figure 3. In the Apo-A 2D RMSD, there are several clear dark squares that appear along the main diagonal line, and they were reflected as low energy areas in the FEL (*i.e.,* AA-S1, AA-S2, AA-S3 and AA-S4). Such squares appear in all three blocks along the diagonal line, therefore, we were able to collect at least one metastable state from each run for Apo-A in the course of 1 *µs* MD simulation. These metastable states seemed to have been visited by the protein in other runs, as apparent with the dark areas shown in off diagonal in the 2D RMSD as well as the trajectory paths in the FEL plot. However, this is mostly true during earlier time in the simulations (*i.e.,* 0-800 ns), and the states visited by the protein at the later time in the simulations (*i.e.,* 800-1000 ns) are quite distinct as evident in the yellow off-diagonal regions of the 2D RMSD. In the FEL plot, while low energy areas appeared, they were not quite separable as clusters, therefore, we were not able to identify clear ensembles for this structure with the current simulation data. Extending simulation time may be needed for detection of clear ensembles in the FEL plot.

**Figure 2:**
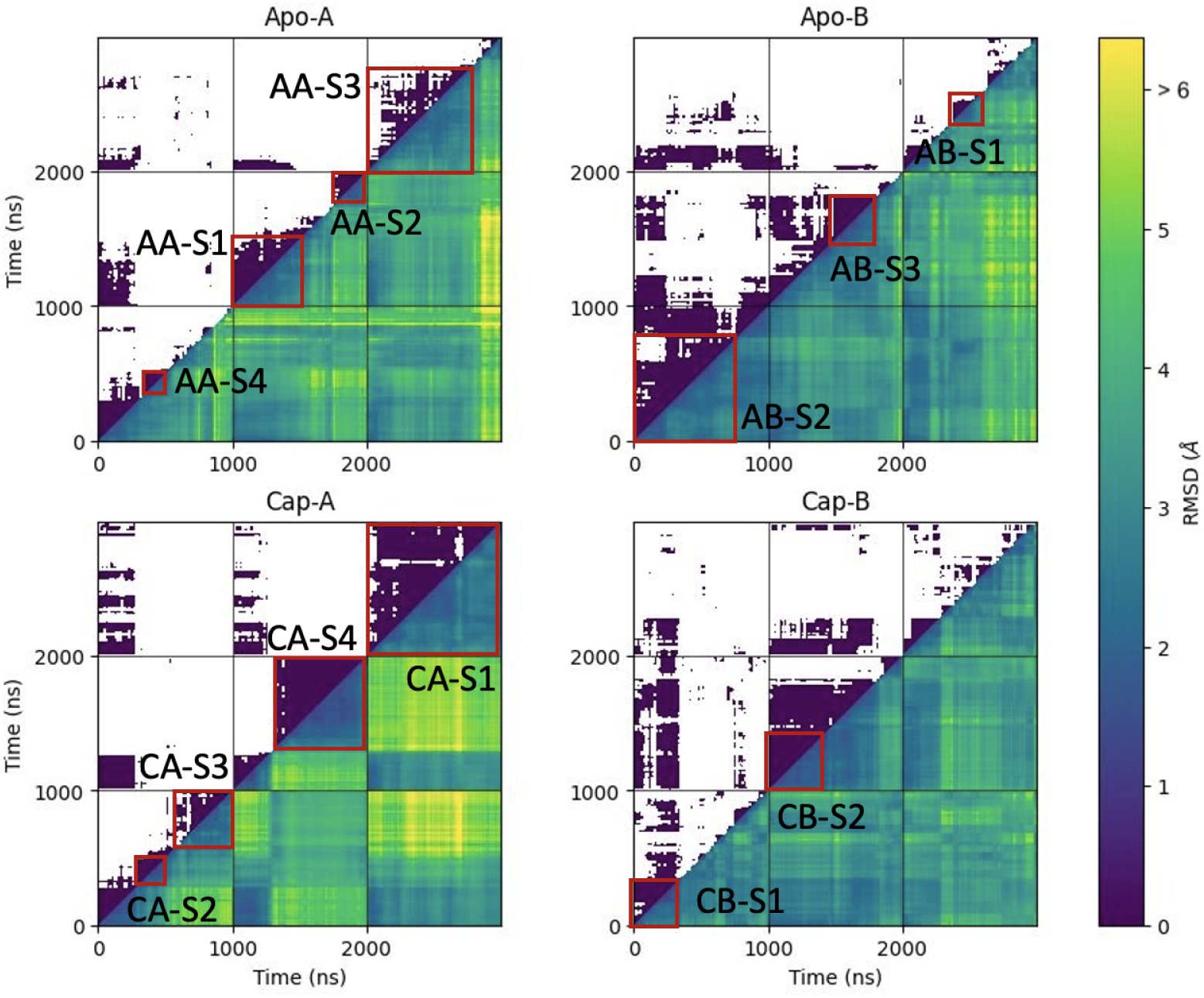
Four 2D RMSD plots show structural dynamics of eFI4E for four different structures. Each 2D RMSD plot shows 2D RMSD of the protein’s concatenated trajectory, where each 0-1000ns, 1000-2000ns and 2000-3000ns in axes correspond to 0-1000ns trajectory of Run 1, Run 2 and Run 3, respectively. The lower triangle is displayed in a heatmap based on the color scale shown on the color bar on the right, and the upper triangle is in binary map with threshold 3.5 Å (*i.e.,* the values less than 3.5 Å is shown in purple and the rest is shown in white.). The low energy areas were identified in FEL, and the corresponding states in 2D RMSD were also highlighted with the red square boxes. Additionally, an arbitrary name was assigned to each identified state so that the states can be distinguished from each other. The names are displayed on right next to the red square boxes in the plot.

**Figure 3:**
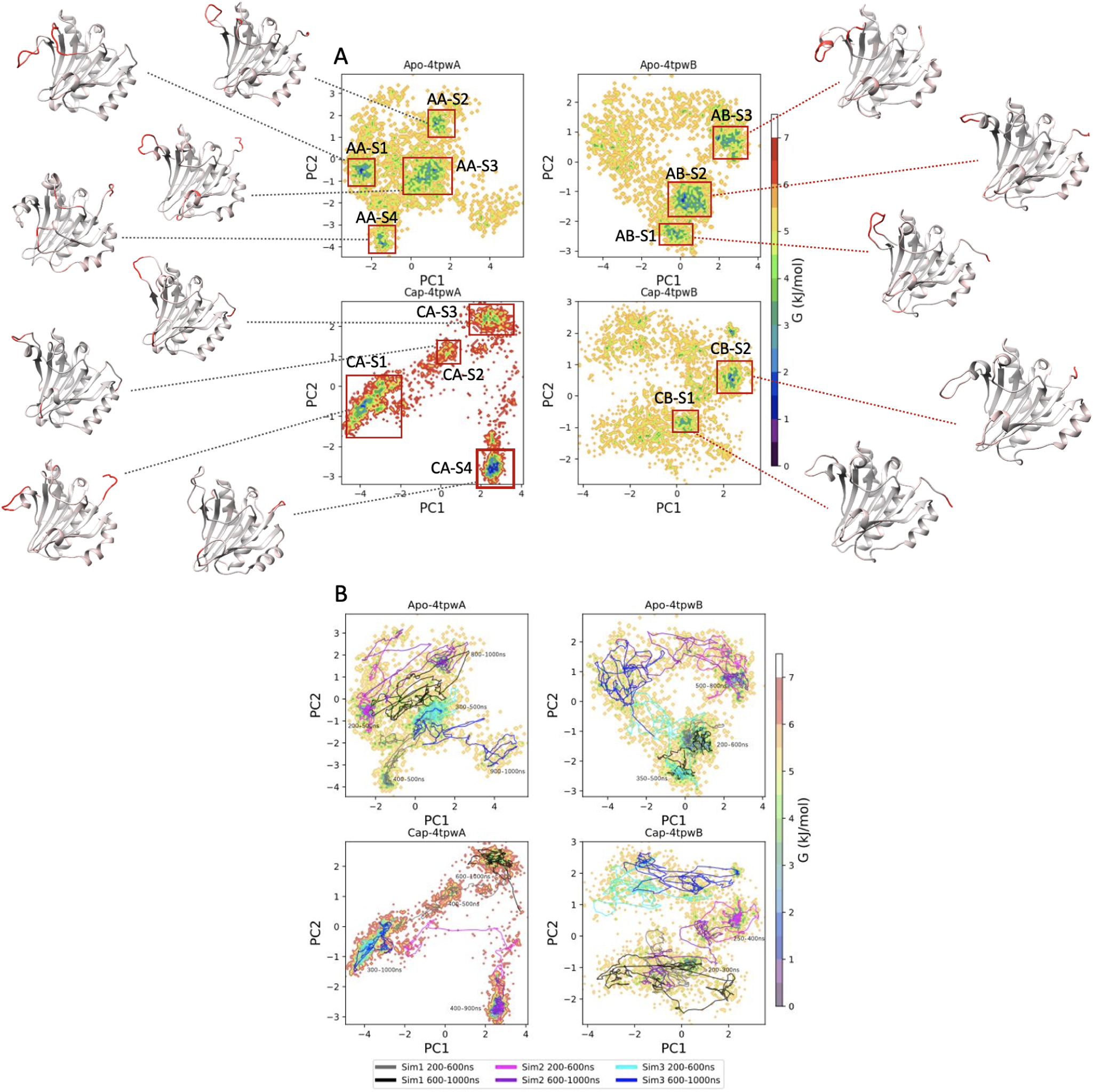
A: Free energy landscape plot with corresponding average eIF4E structures for low energy regions. B: Free energy landscape plot with line paths. The heatmap is scaled based on the color bar on the right, which describes the approximated free energy based on the Boltzmann ensemble. The line paths describes an individual run with two distinct time period, where 200–600 ns - dimgray, 600-100 ns - black for Run 1, 200-600ns - magenta, 600-1000 ns - darkviolet for Run 2, and 200-600 ns - cyan, 600-1000 ns - blue for Run 3.The clusters in FEL that have exhibited low free energies were highlighted with red square boxes. For each structure, representative states were named based on the structure so that the states can be distinguished from each other, and the names were also displayed on both plots right next to the red square boxes. The average eIF4E structures are presented with B-factor color mapping, where red indicates the region has gone through large structural change and gray indicates no structural change at the region with respect to the starting MD structure.

For Apo-B, we have observed a few clear dark squares along the diagonal line in 2D RMSD and they are reflected as low energy regions in the corresponding FEL plot (*i.e.,* AB-S1, AB-S2 and AB-S3). These states are sampled from each of the three runs. The off-diagonal regions in 2D RMSD tend to be darker during earlier simulation time for all three runs, while it got much lighter for the later simulation time for Run 3. This indicates that all three simulations potentially started with a similar state (not shown in FEL as the first 200ns of trajectories are truncated) while during the later simulation time in Run 3 (*i.e.,* 600-1000ns), the protein has visited the states that were not observed in Run 1 and Run 2. Those states that are visited by the protein in later simulation time in Run 3 are also not quite metastable states, evidenced by the lack of dark square(s) on the top right area of the 2D RMSD. In fact, the top left region in FEL where no clusters are formed is mostly dominated by the last half trajectory from Run 3 (*i.e.,* blue line), while other low energy regions in FEL have mixed paths from different runs. Again, similar to the Apo-A, for Apo-B while we were able to sample some metastable states for this structure, the low energy regions appeared in FEL were not clearly separable as clusters.

In Cap-A, four clear squares are formed along the diagonal line (*i.e.,* CA-S1, CA-S2, CA-S3 and CA-S4), at least one for each run. Therefore, at least one metastable state was collected from each run for this structure. Just as the first two protein structure cases, all three runs shared similar states during the early stage of the simulation (0-200 ns) while the states sampled during the later time of the simulations were quite distinct as evident from the yellow areas in the off-diagonal region in the 2D RMSD. As it is expected from the clear separation between the dark squares and the rest of the yellow regions appeared in the 2D RMSD, clear clusters were formed for each state in the FEL. As each states were clearly separated, the protein seems to have gone through a clear transition from one state to another in this structure, which is a different behavior observed in Apo-A and Apo-B cases. For the Cap-B, only one clear dark square appeared along the diagonal line for Run 1 and Run 2 (*i.e.,* CB-S1 and CB-S2), and for Run 3 there was no clear dark square being formed except for during the first 200ns. In the off-diagonal areas, no dark regions appeared except for the first 200 ns of each run just like we observed in the other three structures, indicating that the protein in all three runs started with a similar state and then diverged to distinct states afterwords. For Run 3, even within the same run, the protein has formed multiple structures that are distinct from each other as apparent from the lack of the formation of dark squares in the diagonal line as well as from the trajectory paths in FEL.

Overall, the combined results from the 2D RMSD and the FEL showed that the protein has manifested some metastable structures in all four structures during the 1 *µs* of MD simulations. However, for most of the structures, those states were not able to be separated from one to another very clearly, which could be an indication of insufficient sampling. Moreover, Cap-A structure behaved differently from the rest of three structures, where the transitions from one state to another for Cap-A were much cleaner compared to the other three structures. Cap-A is the 4EGI-1 removed eIF4E/^7^mGTP complex structure. In our past studies,^35,47^ 4EGI-1 has been identified as a promising inhibitor for eIF4E/eIF4G interaction, however, the effect of the 4EGI-1 on eIF4E/^7^mGTP interaction has not been investigated. We hypothesize that the difference we observed in the protein state transitions between Cap-A and the other three structures might be explained with the effect of 4EGI-1 on eIF4E/^7^mGTP interaction. In other words, 4EGI-1 might have stabilized eIF4E/^7^mGTP interaction in 4TPW Chain A structure, and the prolonged effect is still being observed even after the 4EGI-1 is removed from the complex. We will investigate this possibility and present the results in the future release.

### Conformational Change of Cap Binding Residues and ***α***1 Helix

The protein C-*α* RMSF was calculated for two distinct time windows (*i.e.,* 400-500 ns and 900-1000 ns) with respect to its starting (*i.e.,* t = 0 ns) structure to study the local structural change of the protein over the course of 1 *µs* of MD simulations and presented in Figure 4. Across all four systems, the RMSF of the residues that are part of the secondary structures such as *α* helices and beta sheets stayed small, meaning that the local structure in the regions didn’t change much from the starting structure. Meanwhile, the residues that are in the flexible loop regions exhibit high values meaning that the local structure in the regions largely changed since the beginning of the MD simulations. We also observed large RMSF values for some of the Apo-A and Cap-A runs for residues 78-83 that were part of the extended *α*1 helix in their original crystal structures. This is consistent with our expectation, as the removal of the 4EGI-1 from the original 4TPW Chain A (*i.e.,* eIF4E/^7^mGTP/4EGI-1 complex) should affect the structure of the region whose change has originally been induced by the presence of the compound. The residues that are involved and play important role in the ^7^mGTP Cap binding, such as W56, W102, E103 and several lysine and arginine residues, also showed large fluctuations in RMSF plot in all four systems. The fluctuations were, however, particularly noticeable in apo structures (*i.e.,* Apo-A and Apo-B). This could simply be due to the absence (or removal) of ^7^mGTP from the Cap-binding site as those residues no longer have the Cap to interact with.

**Figure 4:**
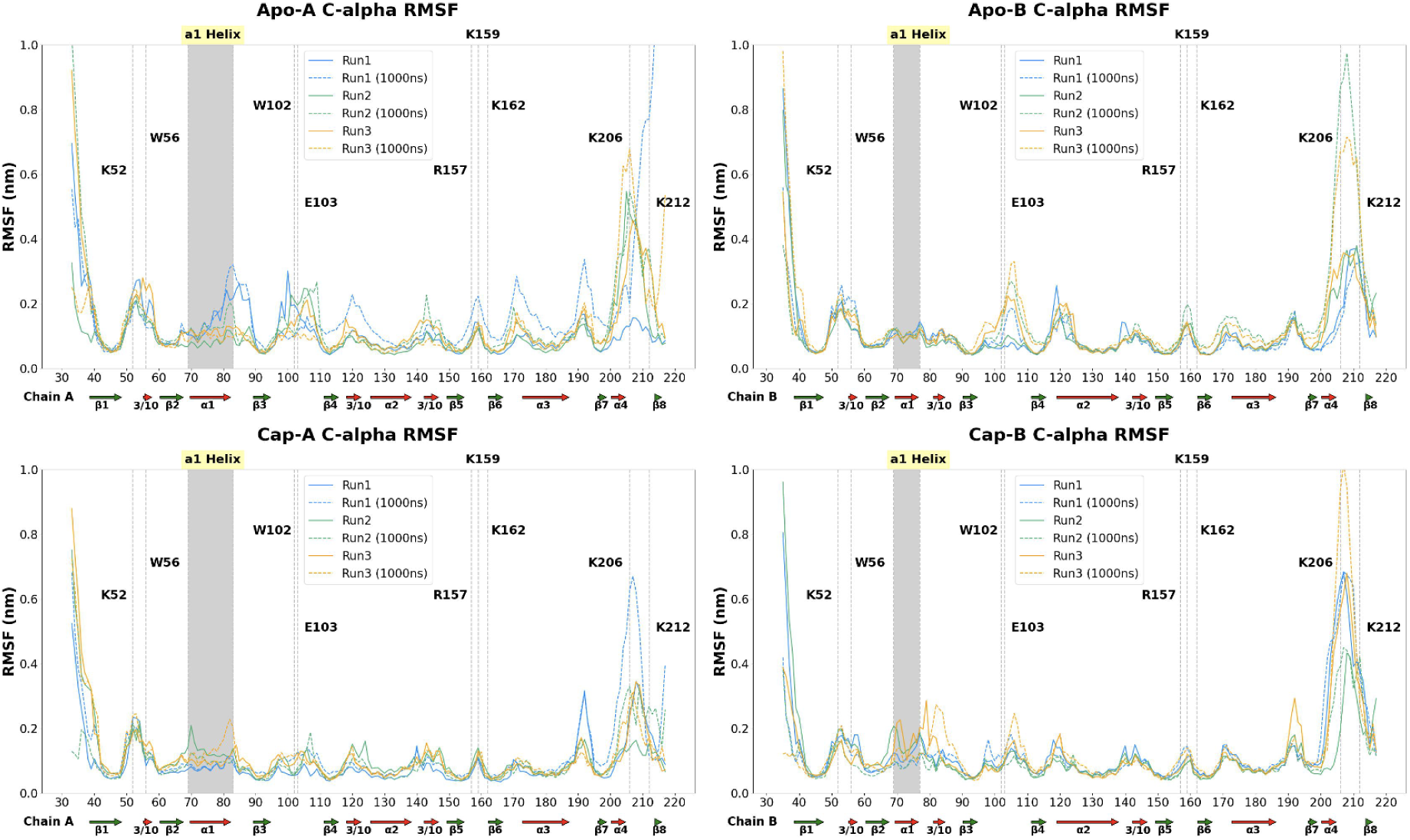
Protein C-*α* RMSF for each eIF4E structure. The x axis is the residue IDs and the y axis is the corresponding RMSF values in nm. RMSF here is calculated for the last 100ns of the 500ns (or 1000ns) trajectory data for each structure. RMSF is calculated based on the difference of the average structure with respect to its 0ns structure. Solid lines are for the 500ns trajectories and dotted lines are for the 1000ns trajectories. Run 1 RMSFs are colored in blue, Run 2 RMSFs are colored in green and Run 3 RMSFs are colored in orange. The key Cap binding residues are annotated, as well as the residues that were part of *α* 1 helix in the original crystal structure were annotated and highlighted with a gray shade. Below the residue IDs, the secondary structure information based on the original crystal structure is displayed.

While RMSF is a useful measurement for quantifying the movements of protein’s local regions, it is often calculated based on a specific time window of a simulated trajectory, and hence lacks the detailed statistics over the course of simulations. For example, in the above generated RMSF plots, we have picked 400-500 ns and 900-1000 ns time windows to calcuate the average fluctuation of each residue. However, the local movements occurred in other time are not reflected in such analysis, hence it is not sufficient to quantify the whole range of residue fluctuations within the collected MD trajectory. Therefore, we have designed and implemented an analysis that can overcome this limitation. We have calculated the distance of each residue from center of mass (*i.e.,* COM) of the protein for every ns of each run and combined the results of three runs into one to calculate the statistics. A box plot was generated for each residue in each structure based on the calculated statistics and presented in Figure 5.

**Figure 5:**
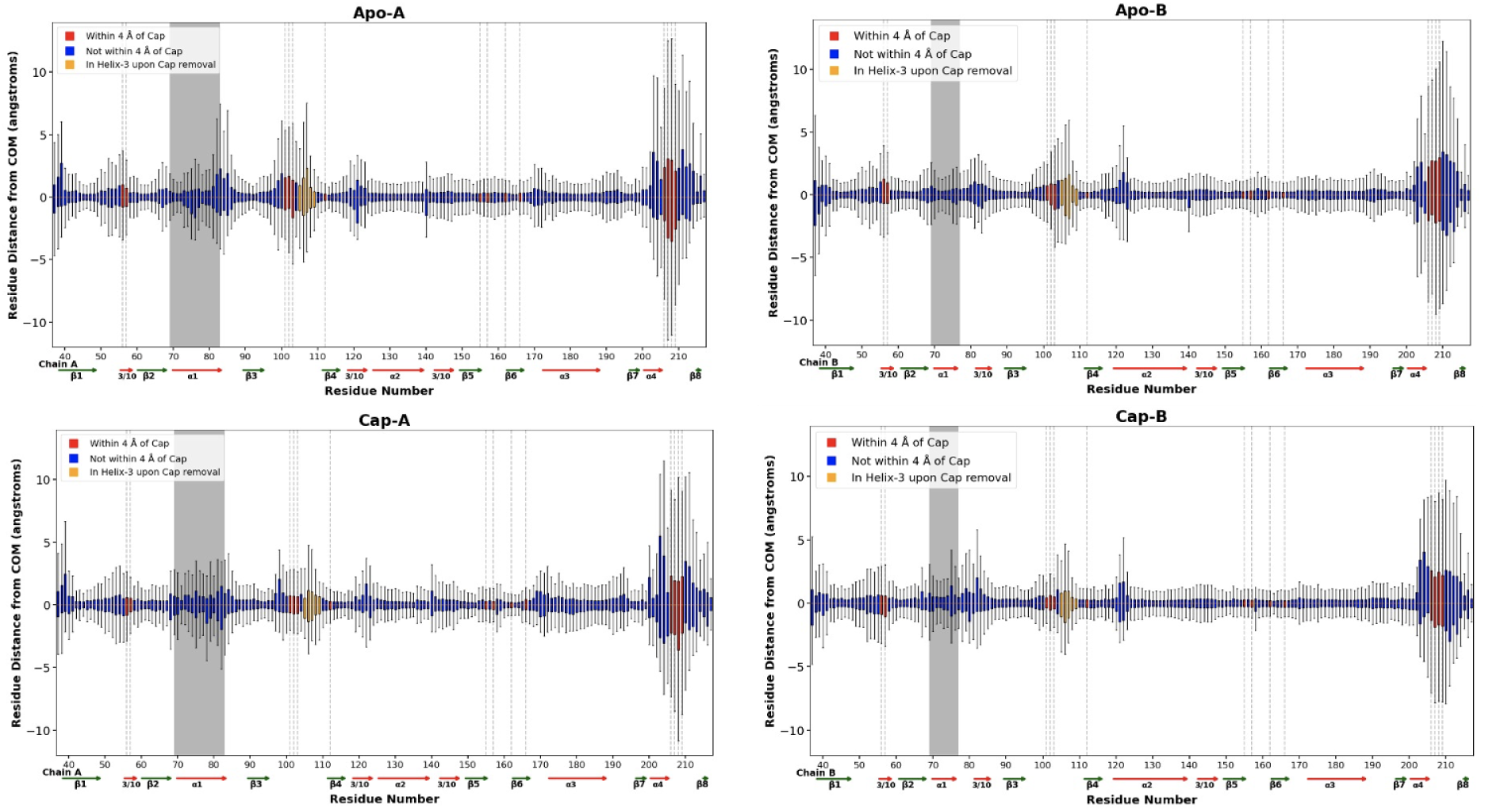
Distance box plots for all the residues in eIF4E presented for each structure in one figure. The concatenated trajectory from all three runs (3*×*1 *µs* = 3*µs*) were used for the calculation. *α* carbons was used as a reference point to take the distance from the COM of the protein. x-axis represents the residue numbers and y-axis represents the distance of the *α* carbon of a residue from the protein COM in Å. The residues that are located within 4 Å of ^7^mGTP in the Cap-bounded eIF4E are colored red, the ones that are located at H3 in 2GPQ apo-eIF4E structure are colored in orange, and the rest are colored blue. Dotted lines are added in the vertical direction to further emphasize the residues that are within 4 Å of ^7^mGTP. Each box is aligned at the mean (*i.e.,* 50th percentile) distance for better visualization.

Overall, the residues located at the *α*-helices and *β*-sheets didn’t fluctuate much with respect to the protein’s COM, while the residues that are located at the loop regions largely fluctuated as evident from the height of the boxes. The key Cap binding residues W102, E103 and W56 also showed large fluctuations while the fluctuations in R157, K159 and K162 were not as large as we could expect from the previous RMSF plots. Other important residues in Cap binding, K52, K162, K206 and K212, have shown large fluctuations as consistent with what we observed in the RMSF results. Based on RMSF and the distance analysis, conformational changes of the local regions in all the protein structures were consistent with what we expected, supporting the validity of our simulations.

### Conformational Change of Helix-3 containing K106 and K108

The mRNA Cap binding pocket does not pre-exist but is induced by ^7^mGTP mRNA Cap through extensive re-arrangement of critical eIF4E residues, namely, W102, E103 and W56. Therefore, one strategy to develop agents for the eIF4E-Cap interaction would be to target the critical residues and prevent the formation of such Cap binding pocket. We might be able to do this by prohibiting the large scale motion of those critical eIF4E residues that are responsible for the pocket formation. Development of bi-valent agents that can simultaneously bind aforementioned residues to a stable structure (e.g. *α* helix containing K106 and K108) would be a potential way to do this. From the previous study,^34^ both K106 and K108 are part of helix-3 that is consisted of residue E105-R109 in apo form (PDB: 2GPQ,^34^ Figure 6 - A). However, in the Cap bound form (PDB: 4TPW Chain B^35^), those residues that were taking a helix structure including K106 and K108 no longer hold a secondary structure and become a part of a flexible loop that connects beta sheets (Figure 6 - B). This is not just for eIF4E/^7^mGTP complex crystal structure (*i.e.,* PDB: 4TPW Chain B), but is also true for eIF4E/^7^mGTP/4EGI-1 complex crystal structure (*i.e.,* PBD: 4TPW Chain A^35^) as presented in Figure 6 - C. This shows that the residues which form the *α* helix in apo-eIF4E go through significant rearrangement during Cap binding. Our MD simulations have captured the intermediate states of the rearrangement due to those residues in the transition from Cap to apo form in both Apo-A and Apo-B cases. The average local structures containing K106 and K108 at two time intervals (*i.e.,* 400-500ns and 900-1000ns) are presented in Figure 6 C-D for Apo-A Run 1 and in Figure 6 E-F for Apo-B as a demonstration. For Apo-A, over the course of 1000ns MD simulations, K106, K108 and the neighboring residues E105, N107 and R109 rearrange the structure and formed an *α* helix at 400-500ns of the trajectory. The NMR Cap free eIF4E structure had E105-R109 as a part of *α* helix, therefore, this *α* helix formed as an intermediate state in Apo-A is identical to the one in NMR apo eIF4E presented in 2GPQ. At 900-1000ns, however, the residues that formed the *α* helix has further rearranged and transitioned back to a loop from an *α* helix. For Apo-B, we observed formations of alpha helix with a subset of residues E105-R109, one with N107-R109 at 400-500ns and another one with E105-N107 at 900-1000ns of the trajectory. These observations in the alpha helix are also reflected as large spikes in the RMSF values as well as the tall box sizes and the large min-max ranges in the distance box plot in residues E105-R109 in Figure 4 and Figure 5, respectively. These were more noticeable in Apo-A and Apo-B cases where the removal of Cap must have influenced the eIF4E structure more than Cap-A and Cap-B cases, hence consistent with our expectations. While the mechanism is not fully uncovered, based on the past studies and from our observation through the MD simulations results, the Cap binding process seems to induce structural rearrangement to the region that contains K106 and K108, causing *α* helix to transform to a loop. Our observations on the available eIF4E structures as well as on our MD simulations suggested that this process is also most likely reversible. These observations might further support our developing approach of designing eIF4E/^7^mGTP inhibitor which works by hinging the movement of the key Cap binding residues W56, W102 and E103 with the *α* helix containing lysine K106 and K108 to prevent the Cap binding pocket formation.

**Figure 6:**
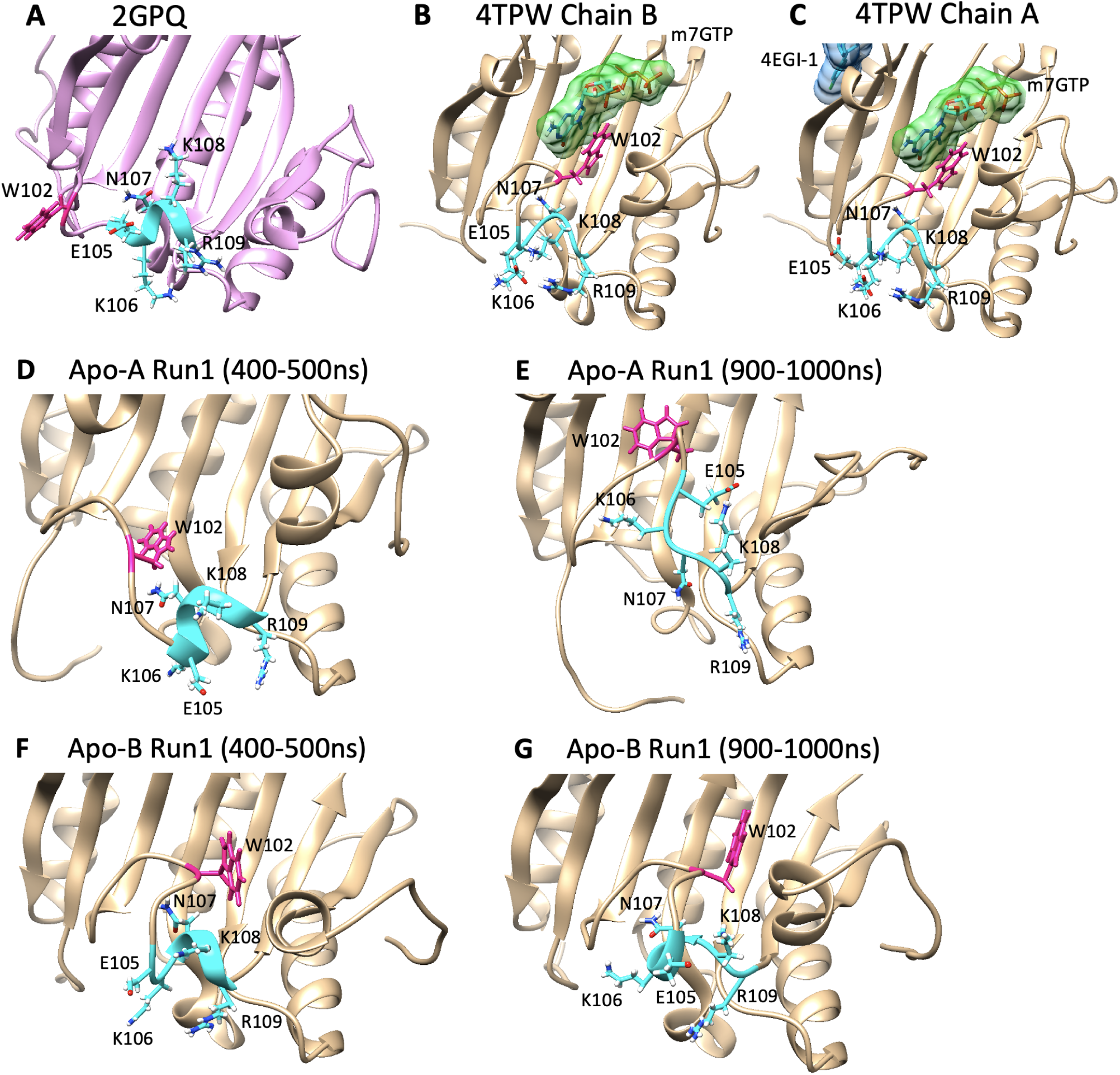
A: 2GPQ, B: 4TPW Chain B, C: 4TPW Chain A, D-E: Average structures of Apo-A Run1 during 400-500ns and 900-1000ns of the MD simulation, F-G: Average structures of Apo-B Run1 during 400-500ns and 900-1000ns of the MD simulation. eIF4E is represented as plum cartoon for 2GPQ and tan cartoon for the rest. ^7^mGTP is shown in cyan stick with solid lime green surface. 4EGI-1 is shown in cyan stick with dodgeblue surface. All the surfaces are shown in 0.7 transparency for better visibility. For E105-R109 and W102, side chains are presented. For E105-R109, both ribbon and the side chains are colored in cyan and for W102, both ribbon and the side chain are colored in deep pink.

## Conclusion

Molecular dynamics (MD) simulations were employed to generate trajectories of the apo and Cap forms of the protein and its complex with mRNA Cap to provide detailed insights into the re-arrangement of critical eIF4E residues, including W102, E103, W56 and the *α*-helix containing key residues such as lysine and arginine. Three independent 1000ns long MD simulations were performed for each Apo-A, Apo-B, Cap-A and Cap-B structure to sample the corresponding protein conformations. We successfully sampled metastable states from the 1 *µs* trajectories for all four structures. Meanwhile, the results from 2D RMSD and FEL showed that we might need additional trajectory data to fully explore the sampling space. Additionally, we have observed the most clear states transition happens in Cap-A. We hypothesize that 4EGI-1 have stabilized the ^7^mGTP-eIF4E interaction such that even after the removal of the ligand the protein remained in a stable state and allowed the protein to go through cleaner state transitions in the simulation. These findings have significant implications for the mechanism of action of 4EGI-1 and design of more potent analogs. RMSF and distance analysis showed the consistent results with the experimental results, supported the validity of our MD simulation. Our observations on apo and Cap eFI4E structures suggested that helix-3 containing K106 and K108 undergoes structural change upon Cap binding, where the region transitions from *α* helix to a loop in the process. This was also true in the case of Cap to apo transition, where we have observed for Apo-A and Apo-B that the region containing E105-R109 transition between *α*-helix and loop through reversible rearrangement during the 1000ns of MD simulations. These observations may further support our developing strategy of designing agents that modulate the movement of critical Cap-binding residues (W56, W102, and E103) along with the alpha helix containing lysine residues (K106 and K108) to disrupt the ^7^mGTP-eIF4E interaction.

As evident in the protein global conformation analysis performed in this paper (e.g. 2D RMSD and FEL) on the collected MD simulation data, triplicates of runs, each 1 *µs*, most likely is not sufficient to capture all conformational states that the each protein structure visits. Two additional runs for each system are currently performed, and the collected data will be presented in the future release to consolidate the findings presented in this paper. Nonetheless, our preliminary results have provided us with the insights of the states eIF4E takes that were previously unknown, and that it could be applied to further develop inhibitors that target the oncogenic protein in future drug design endeavors.

In this paper, we have studied the metastable structures that eFI4E takes in the transition from Cap bounded form to apo form (*i.e.,* Apo-A and Apo-B) as well as Cap bounded form to Cap unbounded form (*i.e.,* Cap-A and Cap-B). However, it is important to further study the apo to Cap transition to deepen the understanding of metastable states the protein takes in the process of Cap binding. Our future work will cover the study of such transition states using an available eIF4E apo structure such as a NMR structure available in PDB (*i.e.,* 2GPQ) to further contribute our knowledge to the future eFI4E drug development.

## Acknowledgement

The project is supported by the SUNY-IBM Consortium Award (PI: Y.D.) and an IBM Faculty Award FP0002468 (PI: Y.D.) through which we accessed the AiMOS supercomputer at Rensselaer Polytechnic Institute and the WSC Cluster at the IBM Thomas J Watson Research Center to carry out all *in silico* experiments reported here. The work was further supported by Brigham Research Institute seed funding to BHA.

## References

(1) Tamarkin-Ben-Harush, A.; Vasseur, J.-J.; Debart, F.; Ulitsky, I.; Dikstein, R. Cap-proximal nucleotides via differential eIF4E binding and alternative promoter usage mediate translational response to energy stress. Elife 2017, 6.

(2) Lazaris-Karatzas, A.; Montine, K. S.; Sonenberg, N. Malignant transformation by a eukaryotic initiation factor subunit that binds to mRNA 5’ cap. Nature 1990, 345, 544–547.

(3) Lazaris-Karatzas, A.;, et al. Ras mediates translation initiation factor 4E-induced malignant transformation. Genes & Development 1992, 6, 1631–1642.

(4) Polunovsky, V. A.;, et al. Translational control of programmed cell death: eukaryotic translation initiation factor 4E blocks apoptosis in growth-factor-restricted fibroblasts with physiologically expressed or deregulated Myc. Molecular and Cellular Biology 1996, 16, 6573–6581.

(5) Polunovsky, V. A.;, et al. Translational control of the antiapoptotic function of Ras. The Journal of Biological Chemistry 2000, 275, 24776–24780.

(6) Petroulakis, E.;, et al. p53-dependent translational control of senescence and transformation via 4E-BPs. Cancer Cell 2009, 16, 439–446.

(7) Siddiqui, N.; Sonenberg, N. Signalling to eIF4E in cancer. Biochemical Society Transactions 2015, 43, 763–772.

(8) Morita, M.;, et al. mTOR Controls Mitochondrial Dynamics and Cell Survival via MTFP1. Molecular Cell 2017, 67, 922–935 e925.

(9) Morita, M.;, et al. mTORC1 controls mitochondrial activity and biogenesis through 4E-BP-dependent translational regulation. Cell Metabolism 2013, 18, 698–711.

(10) Banko, J. L.; Hou, L.; Poulin, F.; Sonenberg, N.; Klann, E. Regulation of eukaryotic initiation factor 4E by converging signaling pathways during metabotropic glutamate receptor-dependent long-term depression. The Journal of Neuroscience 2006, 26, 2167–2173.

(11) Hooshmandi, M.;, et al. 4E-BP2-dependent translation in cerebellar Purkinje cells controls spatial memory but not autism-like behaviors. Cell Reports 2021, 35, 109036.

(12) Hooshmandi, M.;, et al. Excitatory neuron-specific suppression of the integrated stress response contributes to autism-related phenotypes in fragile X syndrome. Neuron 2023, 111, 3028–3040 e3026.

(13) Wiebe, S.;, et al. Cell-type-specific translational control of spatial working memory by the cap-binding protein 4EHP. Molecular Brain 2023, 16, 9.

(14) Sharma, V.;, et al. mRNA translation in astrocytes controls hippocampal long-term synaptic plasticity and memory. Proceedings of the National Academy of Sciences of the United States of America 2023, 120, e2308671120.

(15) Le Bacquer, O.;, et al. 4E-BP1 and 4E-BP2 double knockout mice are protected from aging-associated sarcopenia. Journal of Cachexia, Sarcopenia and Muscle 2019, 10, 696–709.

(16) Crombie, E. M.;, et al. Activation of eIF4E-binding-protein-1 rescues mTORC1-induced sarcopenia by expanding lysosomal degradation capacity. *Journal of Cachexia*, Sarcopenia and Muscle 2023, 14, 198–213.

(17) Avdulov, S.;, et al. Activation of translation complex eIF4F is essential for the genesis and maintenance of the malignant phenotype in human mammary epithelial cells. Cancer Cell 2004, 5, 553–563.

(18) Avdulov, S.;, et al. eIF4E threshold levels differ in governing normal and neoplastic expansion of mammary stem and luminal progenitor cells. Cancer Research 2015, 75, 687–697.

(19) Le Bacquer, O.;, et al. Elevated sensitivity to diet-induced obesity and insulin resistance in mice lacking 4E-BP1 and 4E-BP2. Journal of Clinical Investigation 2007, 117, 387–396.

(20) Rui, L. A link between protein translation and body weight. Journal of Clinical Investigation 2007, 117, 310–313.

(21) Gkogkas, C. G.; Sonenberg, N. Translational control and autism-like behaviors. Cellular Logistics 2013, 3, e24551.

(22) Gkogkas, C. G.;, et al. Autism-related deficits via dysregulated eIF4E-dependent translational control. Nature 2013, 493, 371–377.

(23) Aguilar-Valles, A.;, et al. Inhibition of Group I Metabotropic Glutamate Receptors Reverses Autistic-Like Phenotypes Caused by Deficiency of the Translation Repressor eIF4E Binding Protein 2. The Journal of Neuroscience 2015, 35, 11125–11132.

(24) Chartier-Harlin, M.-C.;, et al. Translation initiator EIF4G1 mutations in familial Parkinson disease. American Journal of Human Genetics 2011, 89, 398–406.

(25) Creus-Muncunill, J.; Ehrlich, M. E. Cell-Autonomous and Non-cell-Autonomous Pathogenic Mechanisms in Huntington’s Disease: Insights from In Vitro and In Vivo Models. Neurotherapeutics 2019, 16, 957–978.

(26) Ghosh, A.;, et al. Alzheimer’s disease-related dysregulation of mRNA translation causes key pathological features with ageing. Translational Psychiatry 2020, 10, 192.

(27) Napoli, I.;, et al. The fragile X syndrome protein represses activity-dependent translation through CYFIP1, a new 4E-BP. Cell 2008, 134, 1042–1054.

(28) Santini, E.;, et al. Reducing eIF4E-eIF4G interactions restores the balance between protein synthesis and actin dynamics in fragile X syndrome model mice. Science Signaling 2017, 10.

(29) Cardenas, E. L.;, et al. Design of Cell-Permeable Inhibitors of Eukaryotic Translation Initiation Factor 4E (eIF4E) for Inhibiting Aberrant Cap-Dependent Translation in Cancer. Journal of Medicinal Chemistry 2023

(30) Yefidoff-Freedman, R.;, et al. 3-substituted indazoles as configurationally locked 4EGI1 mimetics and inhibitors of the eIF4E/eIF4G interaction. Chembiochem 2014, 15, 595–611.

(31) Fischer, P. D.;, et al. A biphenyl inhibitor of eIF4E targeting an internal binding site enables the design of cell-permeable PROTAC-degraders. European Journal of Medicinal Chemistry 2021, 219, 113435.

(32) Wan, X.;, et al. Discovery of Lysine-Targeted eIF4E Inhibitors through Covalent Docking. Journal of the American Chemical Society 2020, 142, 4960–4964.

(33) Takrouri, K.;, et al. Structure-activity relationship study of 4EGI-1, small molecule eIF4E/eIF4G protein-protein interaction inhibitors. European Journal of Medicinal Chemistry 2014, 77, 361–377.

(34) Volpon, L.; Osborne, M. J.; Topisirovic, I.; Siddiqui, N.; Borden, K. L. Cap-free structure of eIF4E suggests a basis for conformational regulation by its ligands. The EMBO Journal 2006, 25, 5138–5149.

(35) Papadopoulos, E. et al. Structure of the eukaryotic translation initiation factor eIF4E in complex with 4EGI-1 reveals an allosteric mechanism for dissociating eIF4G. Proceedings of the National Academy of Sciences 2014, 111, E3187 –E3195.

(36) Jo, S.; Kim, T.; Iyer, V. G.; Im, W. CHARMM-GUI: a web-based graphical user interface for CHARMM. Journal of computational chemistry 2008, 29, 1859–1865.

(37) Guvench, O.; Mallajosyula, S. S.; Raman, E. P.; Hatcher, E.; Vanommeslaeghe, K.; Foster, T. J.; Jamison, F. W.; MacKerell Jr, A. D. CHARMM additive all-atom force field for carbohydrate derivatives and its utility in polysaccharide and carbohydrate– protein modeling. Journal of chemical theory and computation 2011, 7, 3162–3180.

(38) Berendsen, H. J.; van der Spoel, D.; van Drunen, R. GROMACS: A message-passing parallel molecular dynamics implementation. Computer physics communications 1995, 91, 43–56.

(39) Lindahl, E.; Hess, B.; Van Der Spoel, D. GROMACS 3.0: a package for molecular simulation and trajectory analysis. Molecular modeling annual 2001, 7, 306–317.

(40) Van Der Spoel, D.; Lindahl, E.; Hess, B.; Groenhof, G.; Mark, A. E.; Berendsen, H. J. GROMACS: fast, flexible, and free. Journal of computational chemistry 2005, 26, 1701–1718.

(41) Abraham, M. J.; Murtola, T.; Schulz, R.; Páll, S.; Smith, J. C.; Hess, B.; Lindahl, E. GROMACS: High performance molecular simulations through multi-level parallelism from laptops to supercomputers. SoftwareX 2015, 1, 19–25.

(42) MacKerell Jr, A. D.; Bashford, D.; Bellott, M.; Dunbrack Jr, R. L.; Evanseck, J. D.; Field, M. J.; Fischer, S.; Gao, J.; Guo, H.; Ha, S.; others All-atom empirical potential for molecular modeling and dynamics studies of proteins. The journal of physical chemistry B 1998, 102, 3586–3616.

(43) Bussi, G.; Donadio, D.; Parrinello, M. Canonical sampling through velocity rescaling. The Journal of chemical physics 2007, 126.

(44) Carretero-González, R.; Kevrekidis, P.; Kevrekidis, I.; Maroudas, D.; Frantzeskakis, D. A Parrinello–Rahman approach to vortex lattices. Physics Letters A 2005, 341, 128–134.

(45) Michaud-Agrawal, N.; Denning, E. J.; Woolf, T. B.; Beckstein, O. MDAnalysis: a toolkit for the analysis of molecular dynamics simulations. Journal of Computational Chemistry 2011, 32, 2319–2327.

(46) Richard J. Gowers; Max Linke; Jonathan Barnoud; Tyler J. E. Reddy; Manuel N. Melo; Sean L. Seyler; Jan DomaÅ„ski; David L. Dotson; SÃ©bastien Buchoux; Ian M. Kenney; Oliver Beckstein MD Analysis: A Python Package for the Rapid Analysis of Molecular Dynamics Simulations. Proceedings of the 15th Python in Science Conference. 2016; pp 98 –105.

(47) Moerke, N. J.; Aktas, H.; Chen, H.; Cantel, S.; Reibarkh, M. Y.; Fahmy, A.; Gross, J. D.; Degterev, A.; Yuan, J.; Chorev, M.; others Small-molecule inhibition of the interaction between the translation initiation factors eIF4E and eIF4G. Cell 2007, 128, 257–267.

